# Strategies and experimental tips for optimized quantitative single-molecule studies of membrane and membrane-associated proteins

**DOI:** 10.1101/2022.12.13.520047

**Authors:** Raffaella Magrassi, Alessandra Picollo, Alberto Diaspro, Francesca Cella Zanacchi

## Abstract

The study of stoichiometry and supra-molecular organization of membrane (and membrane-associated) proteins plays a key role in understanding membrane structure and function. Single-molecule localization techniques (SML), besides providing imaging at unprecedented resolution, also offer quantitative tools such as stepwise photobleaching (SP) experiments and quantitative single-molecule localization (qSMLM). SML is becoming widely present in imaging core facilities but addressing biological problems by molecular counting experiments still remains not straightforward since experimental approaches for sample preparation require particular attention.

We will focus on the experimental aspects that may prevent successful quantitative SML experiments of membrane-associated proteins.

Depending on the specific experiment, to avoid artifacts and to miscount, fine-tuning of the expression levels and proper staining procedures are required, as well as optimized protocols and controls for counting.

The work aims to highlight the crucial aspects that must be faced when quantitative single-molecule experiments are performed, helping to match the gap between sample preparation and the application of quantitative fluorescence microscopy techniques.

## 1. Introduction

Cellular functions are regulated by membrane proteins that form supra-molecular complexes with the often unknown structural organization. Within this context, determining these complexes’ dynamics and organization is necessary to comprehend biological functions. A variety of hetero-oligomer species plays a relevant role, and monomers, dimers, or tetramers organize themselves, interacting and forming complex structures. Fluorescence microscopy and single molecule techniques, in particular, represent the perfect tool for studying the supra-molecular organization of membrane complexes. SMLM, based on fluorescence localization microscopy (PALM)[1, 2], stochastic optical reconstruction microscopy (STORM)[3, 4], or (PAINT)[5] provides intrinsic information at the molecular scale thanks to the ability to separately record the emission from spatially separated molecules. SMLM techniques have been proposed to study membrane molecular structures, such as membrane receptors and ionic channels, and to understand their molecular mechanisms[6–9]. In particular stepwise photobleaching (SP)[7, 10–12] and quantitative singlemolecule localization microscopy (qSMLM)[13–19] have recently emerged as powerful tools for quantitative insights into membrane proteins stoichiometry and organization[7, 8, 10, 20, 21].

In SP experiments, molecules are identified by measuring the variations of the intensity traces in the image hailing from single spots. Molecules are subsequently photobleached, and the intensity drops recognize the photobleaching events. Measuring the number of downward steps provides the molecules counting[7, 8, 12]. In principle, the number of molecules of interest may be easily extracted by the number of intensity steps observed when assuming a one-to-one ratio between the fluorophores and proteins of interest. Although several advances have been recently made in the SP analysis[22], also using machine learning and bayesian approaches[23–26], still the robustness is often limited to a finite number of steps. SP optimally works in identifying dimers, trimers, and tetramers, but the accuracy decreases when the molecular complexity increases or multiple interacting proteins are recruited in a close region. To this end, quantitative single-molecule localization microscopy (qSMLM) recently emerged as a nicely working tool for quantifying membrane proteins at the cellular level. SMLM requires proper fluorescence labeling of the membrane target proteins by adding fluorescent tags or labeling with specific antibodies conjugated with fluorophores.

Unfortunately, the staining procedures may cause over-counting (due to the uncontrolled number of dyes attached to the proteins or over-expression of genetically encoded tags) or under-counting (due to incomplete and partial labeling efficiency or missing due to incomplete maturation). In this context, fluorescent labeling has to follow optimized procedures to control the fluorophore-tag ratio/stoichiometry and to minimize miscounting errors.

Spatial clustering methods[27] or pair correlation[28] provide further counting accuracy in protein organization studies. Furthermore, several calibration system[13, 14, 22, 28–31] were recently proven to offer copy-number estimation, reducing and avoiding over- and under-estimation errors.

Although their application to membrane proteins may seem straightforward, these techniques often present practical hurdles to prevent precise and accurate protein quantification. The sample preparation procedures should then be optimized to match the sample requirements and minimize miscounting errors. CRISPR/Cas9 is the election procedure for live cell imaging. It provides a 1:1 ratio between the fluorescent tag and the protein of interest since it allows FP sequence insertion in the POI native sequence. This leads to a fusion protein expression closer to the endogenous one[32]. Therefore, SM acquisitions are not affected by protein overexpression derived from transient transfections[33–36] when CRISPR/Cas9 approach is used.

However, obtaining a CRISPR/Cas9 cell line requires effort in terms of materials, work, and time. Furthermore, transient transfection often remains popular and widely employed for preliminary studies of membrane protein assemblies. Given the comparison of costs and benefits, CRISPR/Cas9 approach is still not routinely present in laboratories, and often widespread procedures are still based on transient transfection with FPs. Eventually, FPs also provide the advantage of having an additional tag that may be exploited for further immunofluorescence labeling. In fact, the FP tag may be used as the primary antibody target when there are no reliable antibodies specific to the protein of interest[37–39]

To make quantitative SML widely applicable, fine-tuning the transfection procedures is required to obtain a suitable expression for SM counting experiments.

## 2. Materials and Methods

### Molecular biology

Plasmids employed in this paper were constructed using standard molecular biology techniques utilizing recombinant polymerase chain reaction. All the constructs were validated by DNA sequencing.

### HEK293 cells

HEK293 cells were cultured in DMEM (Gibco-Invitrogen) supplemented with 10% heat-inactivated fetal bovine serum (FBS Invitrogen-Thermo, 2mM glutamine (Gibco-Invitrogen), and 100U/mL penicillin/streptomycin (Gibco-Invitrogen). Cells were passed using DPBS (Gibco-Invitrogen) and trypsin-EDTA (0.25%, Gibco-Invitrogen). HEK293 cells were grown in a humidified tissue culture incubator at 37°C and 5% CO2. Cells where fixed in 1.5% PFA for 7 minutes at room temperature and rinsed 3 times in PBS.

### Expression in HEK293 cells

Transient co-transfection was performed with our construct composed of pcDNA3.1 plasmid encoding for Barttin with m-Cherry tagged at the C-terminal tail and the pcDNA3.1 plasmid encoding for ClC-Ka[40]. HEK-293 cells were seeded in Labtek chambers (#1.5 glass-bottom) one day in advance and then transfected with Effectene Transfection Reagent (QIAGEN) according to the manufacturer’s protocol.

### Expression in *Xenopus oocytes*

After linearization, complementary RNAs (cRNA) were transcribed using the mMessage mMachine SP6 kit (Ambion, Waltham, MA). We produced cRNA from constructs pTLN-barttin-m-Cherry, home made by cloning Barttin with m-Cherry tagged at the C-terminal tail and pTLN h-ClC-Kac cloned by Lorenz, C. et al[41].

ClC-Ka and Barttin-mCherry cRNA were co-injected in 1:2 ratio.

### Oocyte injection

Oocytes were harvested from *Xenopus laevis* frogs as described by Carsten A. et al.[42].

Oocytes were enzymatically defolliculated by 1 hr treatment with collagenase type I A (Sigma Aldrich) in a solution containing (in mM) 90 NaCl, 2 KCl, 1 MgCl2, and 10 Hepes at pH 7.5 with gentle shaking at room temperature. A varying amount of cRNA nanoliters were injected with a microinjector (Nanoject II, Drummond Scientific, Broomall, PA). Injected oocytes were maintained at 18°C in modified Barth’s solution containing 88 mM NaCl, 1 mM KCl, 0.41 mM CaCl2, 0.82 mM MgSO4, 0.33 mM Ca(NO3)2, 2.4 mM NaHCO3, 10 mM Hepes and 10 mg/mL of gentamicin at pH 7.4. One to three days after injection, voltage-clamp measurements were performed as SR experiments.

### Imaging protocol and analysis

Images were acquired with a commercial N-STORM TIRF microscope (Nikon Instruments), equipped with an oil immersion objective (CFI Apo TIRF 100x, NA 1.49). The Nikon Perfect Focus System was applied during the entire recording process. The fluorescence signal emitted was spectrally selected by the four colors dichroic mirrors (ZET405/488/561/647; Chroma) and filtered by a multiband pass filter (ZT405/488/561/647; Chroma). The system is equipped with an EMCCD camera (Andor iXon DU-897, Andor Technologies) and 3 laser lines: 405nm (100 mW laser, Coherent CUBE), 561nm (1200 mW laser, Coherent Sapphire), 647nm (300 mW laser; MPB Communications).

- *Stepwise Photobleaching experiments*: For stepwise photobleaching experiments, an excitation intensity of ~0.1 kW/cm2 @ 647 nm and of ~0.15 kW/cm2 @ 561nm were used. Imaging is performed in continuous mode until all the signal is photobleached.
- *qSTORM acquisitions*: an excitation intensity of ~0.9 kW/cm2 for the 647 nm and an activation intensity <0.25 W/cm2 were set. Using highly inclined illumination, we acquired 20,000 frames for each super-resolution image at a 33 Hz frame rate. An imaging cycle contains one activating light pulse (405 nm) alternated with three readout frames (647 nm). The duration of the acquisition was the same in all experiments. Samples were embedded in GLOX based imaging buffer[43] as oxygen scavenging system (40 mg/mL-1 catalase (Sigma-Aldrich), 0.5 mg/mL-1 glucose oxidase, 10% glucose in PBS) and MEA 10mM (cysteamine MEA (#30070-50G; Sigma-Aldrich) in 360mM Tris-HCl).

### STORM analysis, cluster analysis and quantitative STORM approach

STORM image reconstruction is performed using custom software (Insight3, kindly provided by Dr. Bo Huang of University of California).

The molecules were identified by Gaussian fit, and a baseline threshold of 600 counts/pixel was set. The final images were obtained by plotting each identified molecule as Gaussian distributions and correcting for drift by cross-correlation methods[44].

Cluster analysis of STORM localization datasets is performed with a distancebased clustering algorithm previously published[45] custom-written in MATLAB (The MathWorks, Natick, MA). The code identifies spatial clusters of localizations following the previously published method[13, 45] setting a proper scaling factor whose value determines the degree of clusters’ segmentation and a minimum number of molecules (20mol/cluster) as a threshold to avoid noise.

The quantification of oligomeric states by qSTORM is performed using a previously developed method[13], based on DNA origamis as calibration tools. Briefly, DNA origamis functionalized with a controlled number of molecules are used as a reference to extract N calibration curves fn, corresponding to the median number of localizations. Specifically, STORM imaging and cluster analysis of selected DNA origami structures is performed on selected chassis identified by the colocalization of the TAMRA and Alexa 647 signal. Then, the number of localizations distribution is computed to extract the set of calibration curves fn. At this point, a general distribution of localizations hailing from an unknown mixture of oligomeric states may be fitted to a linear combination of calibration distributions fn. After clustering, the protein copy number quantification is performed on chassis functionalized with 7 handles by fitting the number of localizations’ histogram to a functional form (containing a linear combination of calibration functions). The stop criterion of the fit is defined by minimizing the objective function that provides the optimal number of functions used for the fit (Nmax). For further information and details on qSTORM procedures, we refer to qSTORM protocols previously published[46].

### DNA origami assembly

12-helix bundle DNA origami structures were prepared following previously published procedures[13, 47]. Chassis were folded starting from p8064 scaffold and the oligonucleotide staple sequences as previously described[13, 46] Briefly, 100 nM scaffold (Tilibit Nanosystems), 600 nM core staples (Life Technologies), 3.6 μM handle staples (IDT), and 9μM TAMRA-labeled fluorophore anti-handles (IDT) and 9μM A647-labeled fluorophore anti-handles were mixed in DNA origami folding buffer (5 mM Tris [pH 8.0], 1 mM EDTA and 16 mM MgCl2). Folding was performed with heating to 80°C and cooling in single-degree increments to 65°C for 75 min, followed by cooling in single-degree increments to 30°C for 17.5 hr. We purified the folded chassis by glycerol gradient sedimentation by centrifugation[47] through a 10-45% glycerol gradient in TBE buffer supplemented with 11 mM MgCl2 for 130 min at 242,704g in a SW50.1 rotor (Beckman) at 4°C and collected in fractions. The evaluation was performed by 2% agarose gel electrophoresis, and the fractions containing well-folded monomeric chassis were collected. Further details on the DNA origami folding protocols we referred to previously published protocols[13, 46, 47].

## 3. Results

Our work focuses on users’ main challenges when studying membrane complexes by quantitative single-molecule super-resolution microscopy.

We highlight the critical experimental steps to tune sample preparation procedures and facilitate single-molecule acquisition and quantification. In particular, we will focus our attention on the challenges related to sample preparation: 1) the choice of the proper nano-template and fluorescent labeling strategy 2) the proper tuning of the fluorescence expression levels for quantitative SMLM; 3) the optimization of other experimental aspects that could impair quantification; 4) the choice of the optimal quantitative single molecule strategy to tackle specific biological problems depending on the different environment.

### 3.1. Choice of the proper nanotemplate and fluorescent labelling strategy depending upon the biological problem

#### 3.1.1. Nanotempates for super-resolution

##### Cell lines

The biological question usually determines the biological system employed (secondary cell line, primary cell line, *Xenopus* oocyte). However, it could be necessary to consider also the optimal performance for SM experiments. For example, *Xenopus* oocytes are the best system for expressing membrane molecular channels due to the near absence of their membrane channels and well-separated fluorescent signal. *Xenopus* oocytes allow for the expression of a few ion channel proteins per area, making them suitable for stepwise photobleaching experiments. Nevertheless, oocytes may be replaced by other biological systems, such as transfected mammalian cells[7, 48, 49] for quantitative super-resolution experiments (qSMLM) profiting from higher fluorescence signal. Cells suitable for transient expression of proteins must have peculiar features such as resistance in case of serum-free cultivation (necessary for some transfection protocols), the possibility to be transfected with different techniques, and good cost-effectiveness. Due to the specific cell phenotypic features (for example, a particular thickness or shape), the most commonly used cell lines for transient transfection are HEK293, HeLa, and CHO cells[50]. For its performance in transfection efficiency, the most suitable cell lines are HEK 293 derived from primary human embryonic kidney cells with fragments of adenoviral DNA. HEK 293 cells could achieve high transfection efficiency (80-90%) in procedures optimized[51, 52] for cell density and ratio of plasmid using FuGene HD (Roche) or Lipofectamine 2000 (Invitrogen). The choice for the transfection approach depends on the cell type employed and the form of nucleic acids to be transfected. Due to the level of criticality of the transfection procedures, it is important to include appropriate controls to refine the experimental outcome[52]. Primary cell cultures may be employed for biological questions of interest in the neuroscience field (i.e. neurons, glia cells, ecc...). In this case, additional care must be addressed to the specific sample requirements.

##### Xenopus oocytes

As mentioned above, *Xenopus* oocytes provide sparse membrane channel localization, making them optimal for stepwise photobleaching experiments (SP). In fact, stepwise photobleaching was proposed for determining the membrane subunit composition of multimeric channels expressed in Xenopus oocytes as a template[7, 10, 21]. This method relies on counting chromophores attached to each multimeric complex subunit. Therefore fluorescent fusion proteins are used to avoid non-specific labeling and RNAs injected in Xenopus oocytes as described by Chen et al.[53].

#### 3.1.2. Labeling

To observe in vivo membrane molecular structures with super-resolution microscopy (SRM) a prerequisite is to set up genetically encoded fluorescent proteins (by joint the target protein and the proper fluorescent tag in the same DNA sequence and inserting it in an expression vector).

During the cloning procedure, for membrane-fused FPs to reach their correct localization, the fluorescent protein insertion (often located in extreme NH2 or COOH terminus) has to consider the protein’s primary sequence, containing the protein targeting sequence encoding for membrane intracellular protein localization[54–56].

Therefore, the chosen fluorescent tag must have the right proprieties suitable for the SRM technique needed for the experiments: for example, a single-molecule-based super-resolution imaging technique uses photoactivatable fluorescent proteins because of their helpful features[57] i.e, high photon budget[57, 58] and on-off switching rate (on-off ratio). The most common photoactivatable proteins in this context are PAGFP, Dendra2, mEos2, PAmCherry, and Dronpa[59–69]. Nevertheless, reporter fluorescent proteins (FP) often have a significant size that could perturb the target protein’s localization and interactions. An approach to overcome this potential issue is to employ chemical tags[70] encoded by a small sequence matching with many fluorescent dyes. Among them, the mainly used Snap-tag covalently reacts with O6–benzilguanine (BG) derivatives that carry fluorescent dyes[71–76] while the Halo-tag proteins[71, 77] covalently react with chloroalkane ligand[77]. Stable compounds generated from chemical-tag reactions with their ligand feature a rapid and predictable stoichiometry[75].

Some SMLM techniques (i.e. STORM) work optimally with synthetic dyes and perform better when labeling is done by immunofluorescence. However, despite the mentioned potential challenges, FPs also provide smart advantages and may perform live cell and in vivo imaging[49]. Indeed when using immunofluorescence membrane proteins are labeled by adding primary and specific secondary antibodies[37–39], the target protein may also be localized by exploiting antibodies against the FP itself and fluorophores optimized for SMLM. This procedure is beneficial when there are no reliable monoclonal primary antibodies for the membrane target protein or to reduce the experimental costs. Still, the indirect immunofluorescence approach will induce low control over the ratio protein of interest/fluorophores, and calibration tools combined with qSMLM may mitigate this issue. However, it has to be considered that each fluorescent staining strategy has peculiar advantages and disadvantages (see **Table1**), and the optimal approach need to be evaluated depending upon the specific biological question.

**Table 1.** Characteristics and advantages of fluorescent labeling strategies for quantitative SML

| Labeling techniques | Advantages | Disadvantages |
| --- | --- | --- |
| **Fusion Proteins for membrane proteins labeling** |  |  |
| Fluorescent proteins | - *Live cell* imaging<br>- Good ratio costs/benefits<br>- Target protein may be localized by exploiting antibodies against the FP itself<br>- FPs may be used as the primary antibody target (when there are no antibodies specific to the protein of interest) | -Fluorescent proteins size could perturb the target protein's localization and interactions<br>- Expression levels has to be tuned for SP experiments<br>-Transient /stable transfection could led to over- or under-counting and calibration Is required |
| Chemical tags (SNAP, HALO, ...) | -*Live cell* imaging<br>- Small size minimize perturbation of protein localization<br>- Predictable stoichiometry and control on the number of fluorophores<br>- Flexibility on the fluorescent dyes used | Expression levels has to be tuned for SP experiments<br>-<br>- Transient /stable transfection could led to over- or under-counting |
| **Immuno-staining for membrane proteins labeling** |  |  |
|  | - Reduces the experimental hurdles/complexity<br>- Optimal synthetic dyes suitable for super-resolution<br>- Fast and simple procedure | - Mainly fixed sample<br>- Post-fixation artefacts impairing counting<br>- Tuning of fixed procedure and time fixation is required<br>-Large size (if smaller compounds i.e. fragments/nanobodies) |

### 3.2. Fine-tuning of the fluorescence expression level

Quantification by SRM experiments requires a well-defined fluorescent molecule concentration and may be impaired by artifacts due to over- or under-expression of targeted proteins. Different aspects may be used for fine-tuning the expression levels, thus reaching optimized levels for quantitative SRM. In particular, the transfection efficiency may depend on the DNA concentration and the time interval after transfection. Furthermore, controls are needed to verify that proteins are correctly processed and located. We will now discuss the leading practices for expressing recombinant proteins in mammalian cells to suggest proper guidelines for preparing suitable samples for quantitative superresolution studies.

For SP experiments, there is a strong need to lower the expression until the sparse image spots regime is reached, preventing the artificial expression modulation from impairing the physiological conditions. In transient transfection, DNA concentration and time after transfection play a relevant role, and we exploited their balance to regulate the expression levels. For a suitable amount of expressed protein, the transfection efficiency must be tuned by modifying the ratio of nucleic acids to transfection reagents[78, 79]. Arnold et al. compared the effect of transfection efficiencies in primary human myoblasts by varying nucleic acid ratios using chemical transfection reagents such as FuGENE 6. It is reported that 2 μg of DNA to 5 μL of FuGENE 6 reagent is the proper combination for the best transfection. In contrast, different DNA concentration to reagent ratios did not increase the transfection efficiency[78]. Other works showed that transfection efficiency of the human gastric adenocarcinoma cell line was not achieved using the highest transfection reagent to DNA ratio volume but by testing a range of combinations conform to the cell type[80]. From the literature, no universal transfection procedure appears, and experimental protocols have to be optimized according to the specific case.

Furthermore, the studies of membrane molecular structures with SRM approaches often involve heterodimers composed of more fused proteins, so their co-transfection must account for each vector concentration and the distinctive monomer features. For example, if they differ considerably, whether target or fluorescent protein length, the amount of each DNA has to be balanced considering the different protein sizes and its reverberation/impact/effect/consequences on transcription. Further, the range time in which membrane fluorescent fusion proteins are expressed plays a role since SRM experiments require a well-defined fluorescence molecule concentration and sparseness. It is crucial to overview the time after transfection and the amount of protein expression before proceeding to downstream experiments. A quantitative commonly used approach is a Real-Time Polymerase Chain Reaction that estimates the expression efficiency after transfection by monitoring mRNA production of the membrane proteins[81–83]. RT-PCR approach could also be employed to verify if mRNAs of co-transfected heterodimer fusion proteins are correctly expressed in equal amounts and in the same time range. In every case, RT-PCR evaluated only the transcription of the membrane protein but not if it is correctly processed and located. For these reasons, after RT-PCR determination of the expression time range, it is essential to perform preliminary experiments with the required SRM technique, further optimizing the expression procedures. For heterodimer investigations, it is necessary to consider the pre-condition that both membrane proteins are correctly expressed and located. This fact often leads to a further adjustment in the time window before measurements are performed.

To show a possible approach for sample preparation optimization, we tested expression after transient transfection of Barttin tagged with mCherry in HEK cells. Barttin is an accessory protein of CLC-K channels necessary for their functional expression and co-localized with ClC-Ks in kidney and cochlea[84, 85]. Mutations in Barttin corresponding genes led to Bartter’s syndrome syndrome that is associated with congenital deafness and renal failure[85]. Experimental evidence showed that Barttin is physically associated with ClC-Ks proteins and increases their membrane expression[86].

We first evaluated different DNA concentrations (ranging from 400 to 100 ng) starting from the above-mentioned previous knowledge. Although a reduction of the DNA concentration may reduce the signal, it has to be considered that below a certain threshold may also impair cellular functionality. For this reason, we decided not to reduce the DNA concentration further below a certain threshold (200 ng), and we calibrated the fluorescence expression reducing the time window after transfection. We tested different time intervals (ranging from 12h to 48h) for controlling the expression levels. We observed that times higher than 30h exhibit high signal (**Fig2A**), thus precluding the possibility of observing intensity traces with stepwise photobleaching. In contrast, timing below 18h still shows a significant cytoplasmic background (**Fig.2B**), probably because proteins are not yet fully recruited and translocated in the membrane. The optimal experimental conditions for SP counting and SMLM correspond to acquisition performed 21h after transfection and are shown in **Fig.2C**.

**Figure 1.**
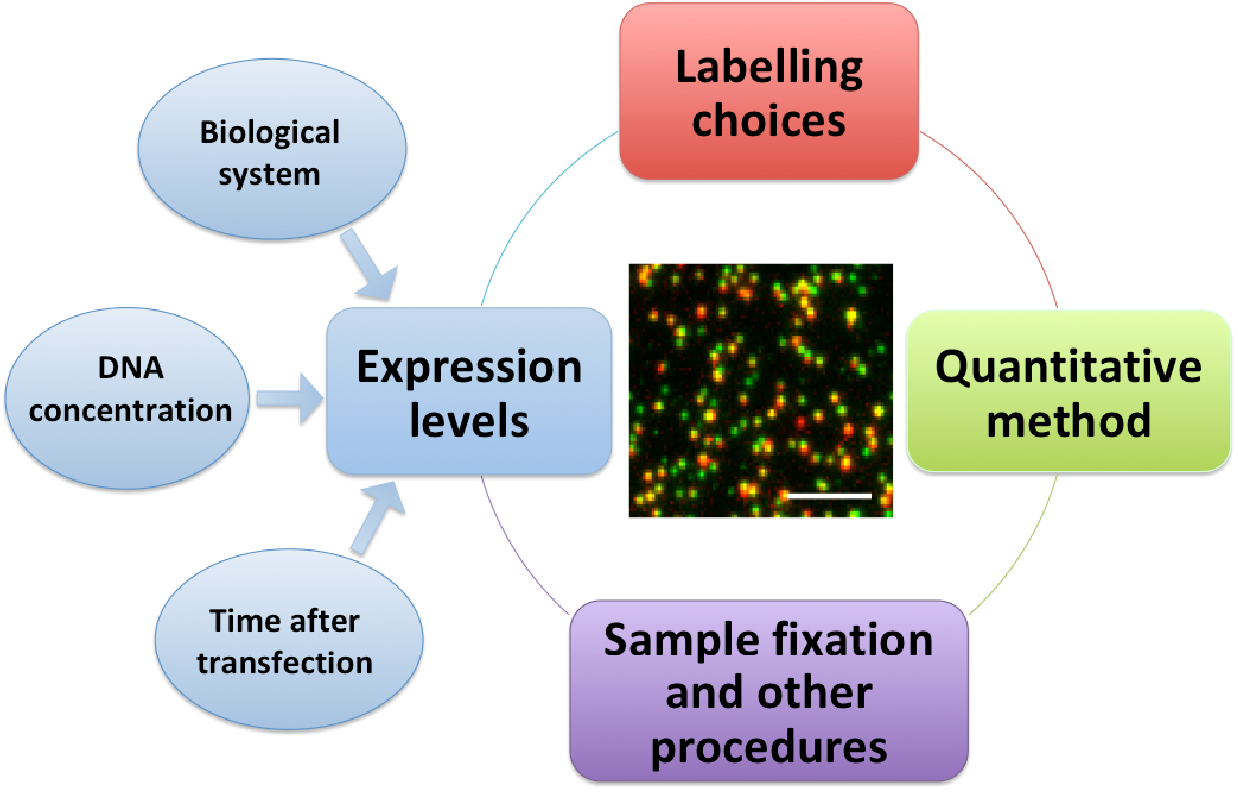
Different challenges for quantitative single-molecule studies of membrane proteins

**Figure 2.**
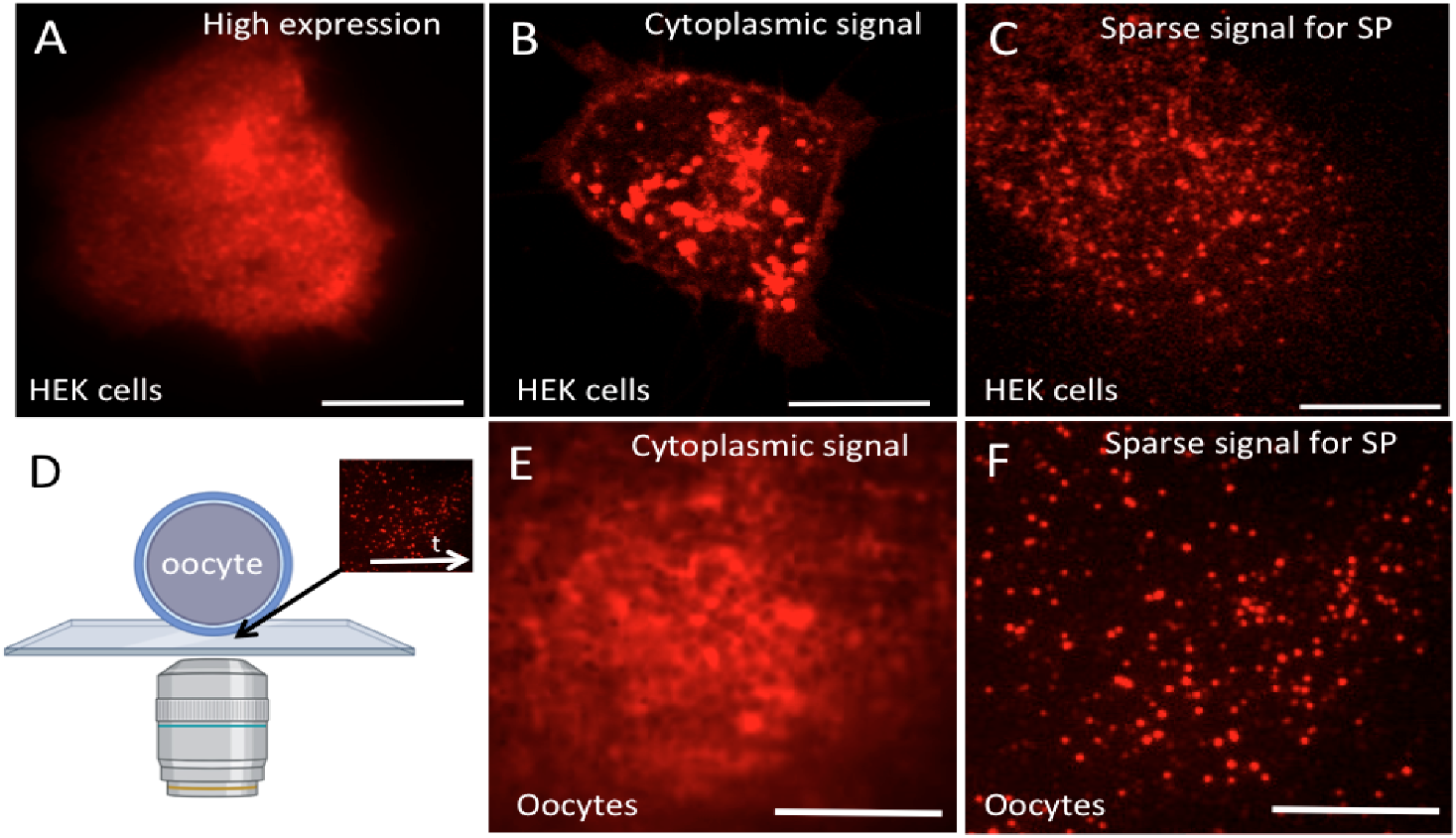
Controlling the expression levels in HEK cells and Xenopus oocytes. Inclined illumination microscopy imaging of HEK cells transiently transfected with Barttin-mCherry (A,B,C) and oocytes (D,E,F). When the time after transfection is longer than 30h the high expression level (A) precludes the possibility to perform SP experiments. Conversely, for time windows shorter than 18h the fluorescence signal is still mainly located in the cytoplasm (B) and proteins are not yet well expressed in the membrane. The correct expression level for stepwise photobleaching experiments is observed (C, Supplementary video 1) for intermediate time intervals (21h). We imaged the Xenopus oocytes membrane (the bottom side adherent to the coverslip) and we monitored Barttin-mCherry by time-lapse experiments (D). SP experiments may be performed only after the cytoplasmic signal (E) disappears and the sparse and low background signal (F) is visible. Scale bars = 10μm

The optimization of the transfection parameters is valid for the specific case, shown as an example. However, a similar concept may also be used to optimize sample preparation procedures for a wide range of proteins of interest.

SRM approaches are often employed in studies that include membrane protein assemblies forming homo or heterodimers. For heterodimer investigations, the experimental complexity increases, and the pre-condition of both proteins correctly located and expressed has to be verified. Furthermore, since proteins are not intrinsically fluorescent but genetically encoded as fusion proteins, it is mandatory before performing downstream experiments to test their functionality. For example, in the case of ion channel complexes, electrophysiology studies have to prove that WT and fluorescent protein-tagged subunits had the same biophysical properties[87].

Heterologous channel expression in Xenopus oocytes depends on the cellular protein processing as synthesis, assembly, post-translational modifications, surface targeting and clustering of the host cell[88]. In addition, in the case of multimeric channel expression, the simultaneous expressions of each component must be obtained by tuning the injected RNA amounts and ratio as a function of the expression time. Indeed fluorescently tagged membrane channels are expressed in oocytes injected with different amounts and ratios of RNAs, and electrophysiological experiments record the corresponding membrane channels’ currents. A proper expression level is reached when similar peaks of the different current magnitudes are found[89].

Since the time after injection plays a significant role, we had to identify the best acquisition settings to visualize a sparse signal optimal for SP experiments in which proteins start reaching the membrane. To this end, we placed the oocyte in the sample holder and performed a time-lapse experiment (**Fig.2D**) to monitor the protein location and identify when the protein translocates from the cytoplasm (**Fig.2E**).

We observed the membrane signal and identified proper conditions for counting experiments by SP, exhibiting a reduced background and avoiding the overlap of undesired cytoplasmic fluorescence (**Fig.2F**).

### 3.3. Optimization and other aspects that could impair quantification

Other experimental procedures, often related to the sample preparation procedure, may be prone to artifacts impairing/impacting the molecular counting capabilities. To this end, particular attention must be given to the protocol optimization and the control experiments required.

Although live cell imaging is preferred, it is not always feasible due to the imaging acquisition process and the limitation on the temporal resolution of quantitative single-molecule experiments. To reduce the experimental hurdles/complexity, users often work on fixed samples rather than living cells for membrane protein quantification.

Although no universal fixation methods have yet been identified as standard best practices for all the samples, we tested different fixation conditions to provide a few general principles for membrane protein counting.

We first considered that organic solvents are not a suitable choice due to the severe loss of membrane integrity induced[90]. On the other side, aldehyde fixation methods[91, 92] perform significantly better, but crosslinking and adduct formation may still be present[90]. Unfortunately, we observed that fixation might interfere with the fluorescence emission signal, inducing a significant drop in the brightness of FPs and a subsequent hurdle in SP counting.

We then tested different aldehyde fixation conditions to optimize the emission signal detected, varying paraformaldehyde concentration, fixation time, and temperature.

We observed that a few adjustments to the commonly suggested fixation conditions (i.e. 3% PFA for 15 minutes at room temperature) helped reduce the detectable signal loss from FPs. Precisely, we followed a fixation protocol (see Methods) that reduces both PFA concentration and fixation time (1.5% PFA for 7 minutes at room temperature).

### 3.4. Choice of the quantitative approach

Proper quantification and analysis of supra-molecular membrane complexes strongly depend on the choice of the quantitative technique used. SP precisely quantifies the oligomeric states of molecular complexes, while qSMLM works optimally for structural information of clustered proteins.

Although the recent developments in increasingly sensitive detectors and bright fluorescent markers improved the accuracy of molecular counting using SP, the number of detectable subunits remains limited (probably less than 8 molecules). Still, despite being the method of choice for studying membrane protein stoichiometry, the maximum number of detectable subunits is not universally determined. We tested the SP performances on a DNA origami model structure (**Fig.3A**) to verify when the intensity traces start to be less robust for revealing the single molecule intensity steps. Twelve-helix structures previously published (Derr, Cella Zanacchi) were functionalized with N molecules of Alexa 647 and 5 molecules of TAMRA (as reference) (**Fig.3A**). To select a signal from the DNA origami nanostructures, avoiding aspecific signal, we imaged by stepwise photobleaching only the spots that exhibit Alexa647 and TAMRA colocalization (**Fig.3B**). We showed that the ability to discriminate intensity steps from the intensity traces is optimal for N=3 Alexa 647 molecules (**Fig.3C**) but it declines when the number of molecules increases (**Fig.3D**).

**Figure 3.**
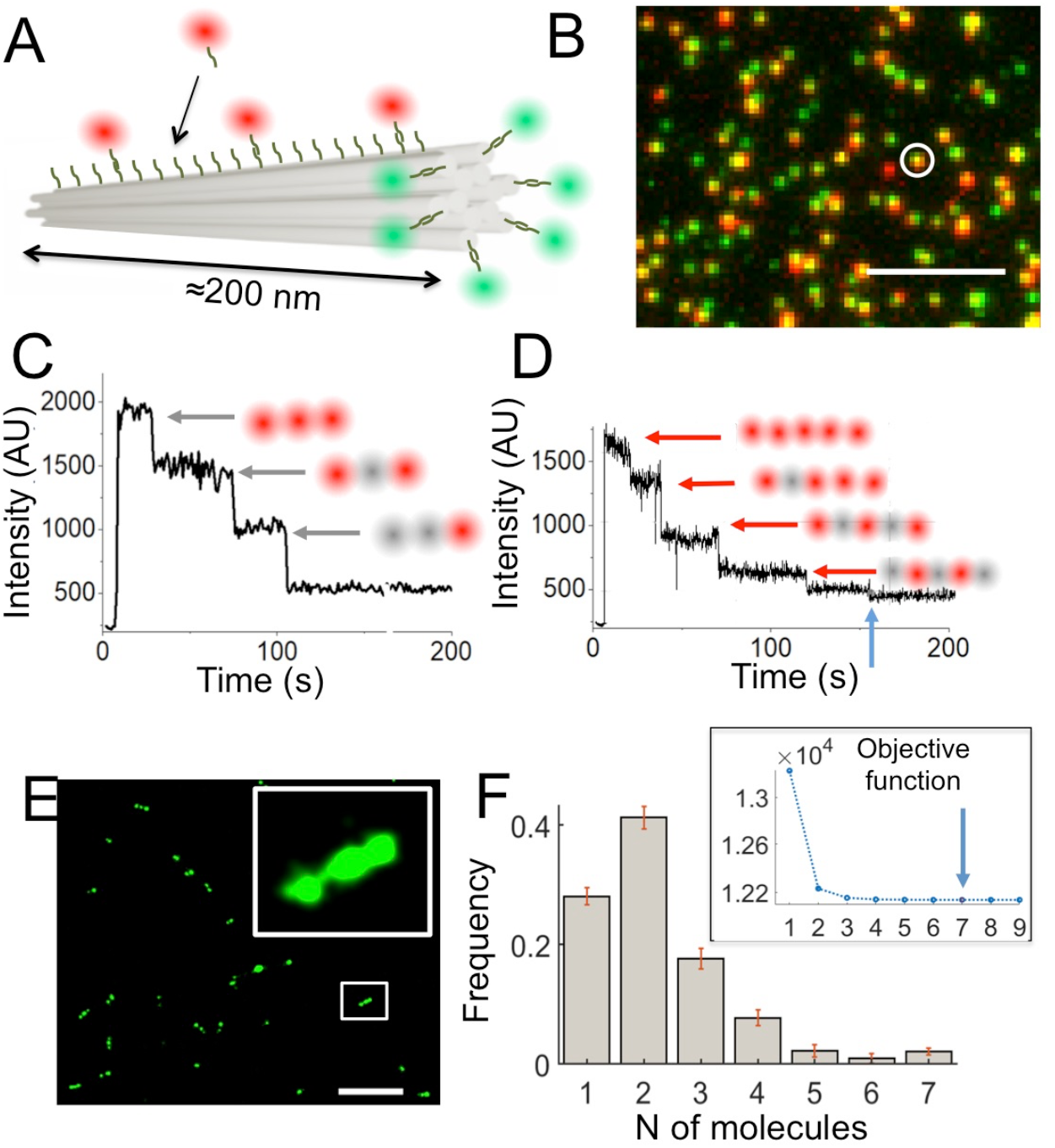
Quantitative Single-Molecule imaging. DNA origamis structures functionalized with different number of fluorophores: 12 helix chassis with N=3 and N=5 Alexa647 fluorophore and 5 TAMRA molecules as reference (A). Widefield imaging shows colocalization of TAMRA and Alexa647 fluorophores on the DNA origamis. Stepwise photobleaching experiments for N=3 (C) and N=5 (D) functionalized Handles on the DNA nanostructure. STORM imaging of DNA origami with N=7 Alexa 647 fluorophores (E) and corresponding oligomeric states quantificationby qSTORM (F). The minimization of the objective function (inset) defines the maximum number of molecules identified in the sample (N=7). Scale bar 1μm (B) and 150nm (E).

Intensity traces show difficulties in evaluating the correct steps’ number from the intensity drop already arising for DNA structures labeled with N=5 Alexa647 molecules (as shown by the barely visible step in **Fig.3D, blu arrow**). Although SP optimally works in identifying dimers, trimers, and tetramers, the accuracy decreases when the molecular complexity increases. In this case, quantitative SMLM may represent a solution. To show the performances of the different quantitative methods, we quantified the DNA origami structures with 7 labeled handles using a quantitative single-molecule method based on STORM microscopy (as previously described[13], see **Methods**). Briefly, we performed STORM imaging (**Fig.3E, zoomed region**) followed by clustering analysis [see **Methods**]. We then used DNA origami as calibration tools (see **Methods**) for extracting the oligomeric states’ (**Fig.3F**) and proteins’ copy numbers from the localizations distribution. The objective function determines the stop criteria for the fitting (N=7, **Fig.3F inset**, blue arrow). This value demonstrates that the oligomeric states calculated by qSTORM well match the effective distribution (DNA origami with 7 handles) and shows the result’s robustness. The widespread distribution of the oligomeric states is mainly due to the expected partial labeling efficiency.

The experiments mentioned above suggest that the results’ accuracy strongly depends on the number of proteins involved and that qSMLM may provide more robust results when the protein copy number increases. Therefore, qSTORM becomes the preferred tool for molecular counting when molecular numbers increase, or several molecules are recruited nearby.

When the membrane protein organization increases, qSTORM may be considered a valid alternative to stepwise photobleaching experiments and different applications require different methods (see **Table 2**).

**Table 2.** Applications of different quantitative single molecule approaches

| Imaging Technique | Oligomeric states of individual molecular complexes | Structural information about individual oligomers | Protein aggregates and clusters | Living systems |
| --- | --- | --- | --- | --- |
| Stepwise Photobleaching | x |  |  | x |
| qSMLM |  | x | x |  |

## 4. Discussion

The work aims to highlight the different aspects required for transforming quantitative SML into an efficient tool for addressing specific biological questions involving membrane (and membrane-associated proteins). Providing practical tips on basic sample preparation procedures optimized for quantitative superresolution microscopy will help make these techniques popular in labs and facilities with limited experience in SML techniques.

Although the expression level may be tuned to facilitate quantitative SMLM, some limitations still remain. Despite several calibration standards being proposed[13, 14, 29–31] a universal best practice has not yet been identified for all the applications. Calibration may be carried out by a) synthetic nanostructures[13, 14, 17, 93] (i.e. DNA origami) or b) with monomeric or multimeric variants of known intracellular proteins/molecules[29–31]. Calibration employing intracellular monomeric proteins may fail accuracy when proteins are organized in densely packed regions. In fact, in the neuroscience context, some biological structures (such as the synaptic environments) present a highly dense environment. The high number of synaptic molecules [94, 95] has to be adequately mimicked by the calibration structures chosen, otherwise the different characteristics may impair a precise quantification. In this scenario, the synthetic structures provide higher flexibility in tuning the number of attached proteins and fluorophores. In this context, smart solutions to tune and increase the number of functionalized handles[96] may extend molecular counting to protein assemblies, improving the mimicking of highly dense protein clusters. This may extend the accuracy of these techniques for synaptic studies[95] that usually involve proteins organized in dense and packed clusters.

## Author Contributions

Conceptualization, F.C.Z., R.M. and A.D.; methodology, F.C.Z., R.M. and A.P.; software, F.C.Z.; formal analysis, R.M., F.C.Z.; investigation, R.M., F.C.Z.; resources, F.C.Z., R.M., A.P. and A.D.; data curation, F.C.Z. and R.M.; writing — original draft F.C.Z., R.M.; F.C.Z., R.M., A.P. and A.D.; writing — review and editing. All authors have read and agreed to the published version of the manuscript.

## Institutional Review Board Statement

The animal study protocol was approved by the Institutional Review Board (or Ethics Committee) of Biophysics Institute, CNR (protocol code 205/2020-PR).

## Acknowledgments

We acknowledge the Nikon Imaging center @ Istituto Italiano di Tecnologia

